# Human serum induces daptomycin tolerance in *Enterococcus faecalis* and viridans group streptococci

**DOI:** 10.1101/2022.10.12.511873

**Authors:** Alicia R. H. Tickle, Elizabeth V. K. Ledger, Andrew M. Edwards

**Affiliations:** MRC Centre for Molecular Bacteriology and Infection, Imperial College London, Armstrong Rd, London, SW7 2AZ, UK; Southmead Hospital, Southmead Road, Westbury-on-Trym, Bristol, Avon, BS10 5NB, UK

**Keywords:** *Enterococcus*, *Streptococcus*, daptomycin, antibiotic tolerance, peptidoglycan

## Abstract

Daptomycin is a membrane-targeting lipopeptide antibiotic used in the treatment of infective endocarditis caused by multidrug-resistant Gram-positive bacteria such as *Staphylococcus aureus*, enterococci and viridans group streptococci. Despite demonstrating excellent *in vitro* activity and a low prevalence of resistant isolates, treatment failure is a significant concern, particularly for enterococcal infection. We have shown recently that human serum triggers daptomycin tolerance in *S. aureus*, but it was not clear if a similar phenotype occurred in other major infective endocarditis pathogens. We found that *Enterococcus faecalis, Streptococcus gordonii* or *Streptococcus mutans* grown under standard laboratory conditions were efficiently killed by daptomycin, whereas bacteria pre-incubated in human serum survived exposure to the antibiotic, with >99% cells remaining viable. Incubation of enterococci or streptococci in serum led to peptidoglycan accumulation, as shown by increased incorporation of the fluorescent D-amino analogue HADA. Inhibition of peptidoglycan accumulation using the antibiotic fosfomycin resulted in a >10-fold reduction in serum-induced daptomycin tolerance, demonstrating the important contribution of the cell wall to the phenotype. We also identified a small contribution to daptomycin tolerance in *E. faecalis* from cardiolipin synthases, although this may reflect the inherent susceptibility of cardiolipin-deficient mutants. In summary, serum-induced daptomycin tolerance is a consistent phenomenon between Gram-positive infective endocarditis pathogens, but it may be mitigated using currently available antibiotic combination therapy.

## Introduction

Infective endocarditis (IE) is a bacterial infection of the cardiac endothelium, primarily affecting the valves. It is relatively rare, affecting 3-10 per 100,000 people annually, but it is fatal if left untreated [1,2]. Despite advancements in medical and surgical treatments for IE, the in-hospital mortality is 15-20%, rising to 40% after one year [1,2]. People at greatest risk of IE are those with pre-existing heart disease, a prosthetic heart valve, cancer, diabetes, the elderly or intravenous drug users [1,2,3].

IE occurs when bacteria enter the blood via medical or dental procedures including toothbrushing [4,5]. Bacteria then colonise the heart valves, particularly when they are damaged or have been replaced by a prosthetic device, triggering the formation of sterile platelet-fibrin microthrombi, to which bacteria can easily adhere [1,2,5,6,7].

Over 80% of cases of IE are caused by Gram-positive cocci, with one cohort study including 2781 participants revealing that *Staphylococcus aureus*, viridans group streptococci (VGS), and enterococci accounted for 31%, 17% and 10% of cases respectively [3]. Leading VGS species include *Streptococcus gordonnii* and *Streptococcus mutans* [8], whilst *Enterococcus faecalis* is the most common enterococcal cause of IE [9].

Host survival depends on several weeks of antibiotic treatment [10], most commonly with a beta-lactam, in combination with gentamicin if an enterococcal species is involved [10]. Even with this duration of antibiotics, surgery is often required to completely eradicate infection and repair the damaged valve [10,11].

Added to the challenges of IE treatment, the emergence of antibiotic resistance requires the use of less efficacious antibiotics. For example, vancomycin, a glycopeptide antibiotic, has long been used to treat infections caused by multidrug-resistant Gram-positive bacteria [12,13]. However, 30% of enterococcal isolates in the USA are resistant, with the Centers for Disease Control and Prevention (CDC) classifying VRE as a serious threat to public health [13]. The rise in multi-drug resistant VGS is also a growing threat [14], with the SENTRY Antimicrobial Surveillance Program estimating that approximately 31% are not fully susceptible to penicillin.

Since its approval for clinical use in 2003, the calcium-dependent cyclic lipopeptide antibiotic daptomycin has become an increasingly used option against multidrug-resistant Gram-positive bacteria, including VRE [15,16,17].

Daptomycin demonstrates excellent *in vitro* bactericidal activity against Gram-positive bacteria via a mechanism that disrupts both the membrane and cell wall biosynthesis [15,16,17,18,19,20,21,22]. Furthermore, while the emergence of resistance to daptomycin has been reported in enterococci and streptococci, it is rare, with over 99% of enterococcal and streptococcal clinical isolates susceptible to the lipopeptide antibiotic [23,24,25]. However, despite the low incidence of resistance to daptomycin and the antibiotic’s potent *in vitro* activity, daptomycin treatment fails to cure 14-24% of patients with enterococcal infection, resulting in poor patient prognoses [26].

The reasons behind the disparity between *in vitro* and *in vivo* efficacies are poorly understood, but there is increasing evidence that the host environment induces stress responses in bacteria that reduces their susceptibility to antibiotics [27]. For example, we have recently shown that human serum induces daptomycin tolerance in *S. aureus* via changes to the cell wall and membrane, mediated in part via the induction of the GraRS two component system by the antimicrobial peptide LL-37 [28]. However, it is unknown whether this phenomenon is specific to *S. aureus* or also occurs in other Gram-positive IE pathogens and therefore, the aim of this work was to determine whether human serum triggered daptomycin tolerance in *E. faecalis* and VGS.

## Methods

### Bacterial strains and growth conditions

Bacterial strains used in this study are shown in Table 1. *E. faecalis* strains JH2-2 [29] and OG1X [30]; *S. gordonii* strain DL1 [31]; and *S. mutans* strain UA159 [32] were grown in 3 ml Todd-Hewitt broth, supplemented with 1% Yeast extract (THY) at 37°C in 5% CO_2_ for 16 hours (h) to stationary phase. For experiments with daptomycin, THY was supplemented with 1.25 mM CaCl_2_.

**Table 1.** Strains used in this study.

| Strain | Relevant characteristics | Reference |
| --- | --- | --- |
| *Enterococcus faecalis* JH2-2 | Fus<sup>R</sup> Rif<sup>R</sup> strain derived from non-haemolytic clinical isolate JH2 | 29 |
| *Enterococcus faecalis* OG1X | Sm<sup>R</sup> derivative of clinical isolate OG1 | 30 |
| *Enterococcus faecalis* OG1RF | Sm<sup>S</sup> derivative of clinical isolate OG1 | 47 |
| OG1RF Δ*cls1* | OG1RF with *cls1* deleted | 39 |
| OG1RF Δ*cls2* | OG1RF with *cls2* deleted | 39 |
| OG1RF Δ*cls1*/Δ*cls2* | OG1RF with *cls1* and *cls2* deleted | 39 |
| OG1RF Δ*mprF2* | OG1RF with *mprF2* deleted | 39 |
| OG1RF Δ*cls1*/Δ*cls2*/Δ*mprF2* | OG1RF with *cls1*, *cls2* and *mprF2* deleted | 39 |
| *Streptococcus gordonii* DL1 |  | 31 |
| *Streptococcus mutans* UA159 |  | 32 |

### Minimum inhibitory concentration (MIC) determination

The MICs of daptomycin were determined using the standard clinical laboratory standards institute broth microdilution technique [33]. In brief, two-fold serial dilutions of antibiotics were made in Müller-Hinton broth supplemented with 1.25 mM CaCl_2_ in 96 well plates before they were inoculated with 1 × 10^5^ CFU ml^-1^ bacteria and incubated statically at 37°C in 5% CO_2_ for 16 h. The lowest concentration at which there was no visible growth was determined to be the MIC.

### Daptomycin survival assay

To prepare THY-grown bacteria, overnight cultures were diluted to 10^7^ CFU ml^-1^ and then incubated statically for 2 h to 10^8^ CFU ml^-1^ at 37°C in 5% CO_2_. To generate serum-adapted cultures, THY-grown bacteria were recovered by centrifugation, resuspended in an equal volume of human serum (Sigma) and incubated statically for 16 h at 37°C in 5% CO_2_. Where appropriate, indicated concentrations of fosfomycin were added during serum-adaptation. Immediately before the assay, THY-grown cultures were centrifuged and resuspended in serum. THY-grown and serum-adapted cultures were incubated with or without daptomycin (80 μg ml^-1^) for 6 h statically for 37°C in 5% CO_2_.

Before and after the 6 h daptomycin exposure, bacteria were enumerated via 10-fold serial dilution in PBS and plating onto THY agar before incubation for 16 h at 37°C in 5% CO_2_. The number of colonies in dilutions which yielded 10 – 100 colonies were counted and used to determine CFU ml^-1^.

### Peptidoglycan accumulation assay

The synthesis of peptidoglycan was determined by measuring the incorporation of the D-alanine analogue HADA [34]. THY-grown and serum-adapted bacteria were generated as described above in the presence of HADA (25 μM) and in the dark to prevent bleaching of the fluorophore before being fixed in 4 % paraformaldehyde.

To quantify HADA incorporation, bacteria were spotted onto microscope slides covered with a thin agarose layer (1.2 % agarose in water) and covered with a cover slip. Phase-contrast and fluorescence images were taken using a Zeiss Axio Imager.A1 microscope coupled to an AxioCam MRm and a 100x objective and processed using the Zen 2012 software (blue edition). HADA was detected using the DAPI filter set. Fluorescence of the cell walls of individual cells was quantified using ImageJ. All images were taken on the same day using identical settings to allow comparisons to be made between samples.

### Statistical analyses

CFU data were log transformed and represented as the geometric mean ± geometric standard deviation of the number of independent experiments indicated in the figure legends. Data were analysed by paired t-test, one-way ANOVA or two-way ANOVA with appropriate multiple comparisons test, as described in the figure legends using GraphPad Prism 8.

## Results

### Human serum induces daptomycin tolerance in *E. faecalis* and representative members of viridans group streptococci

To test whether serum induces daptomycin tolerance in IE pathogens besides *S. aureus* we employed a panel of four different strains: *E. faecalis* JH2-2, *E. faecalis* OG1X, *S. gordonii* DL1 and *S. mutans* UA159. Bacteria were grown in THY broth (THY-grown) to represent *in vitro* conditions or incubated in human serum (serum-adapted) to mimic conditions found during the host. Bacteria were then exposed to daptomycin at 80 μg ml^-1^, or left untreated, and bacterial survival measured via determination of CFU counts after 6 h. Exposure to daptomycin was carried out in serum to control for binding of daptomycin by plasma proteins and to keep the calcium concentrations consistent [35]. This concentration of antibiotic was chosen as it corresponds to the peak serum concentrations during therapy [36] and is well in excess of the daptomycin MIC of each of the strains (Table 2).

**Table 2.** Daptomycin MIC of strains used in this study.

| Strain | Daptomycin MIC (µg ml<sup>-1</sup>) |
| --- | --- |
| *E. faecalis* JH2-2 | 0.5 |
| *E. faecalis* OG1X | 0.5 |
| *S. gordonii* DL1 | 2 |
| *S. mutans* UA159 | 0.25 |

As expected from previous reports, daptomycin was rapidly bactericidal against THY-grown cultures of all four strains, with ∼40-6000-fold reductions in CFU counts of bacteria surviving exposure to 80 μg ml^-1^ daptomycin after 6 h [37]. By contrast, > 99 % of all four strains of serum-adapted bacteria survived 6 h exposure to 80 μg ml^-1^ daptomycin (Fig. 1). Therefore, as seen previously for *S. aureus*, serum adaptation of *E. faecalis* and VGS resulted in high levels of daptomycin tolerance [28].

**Figure 1.**
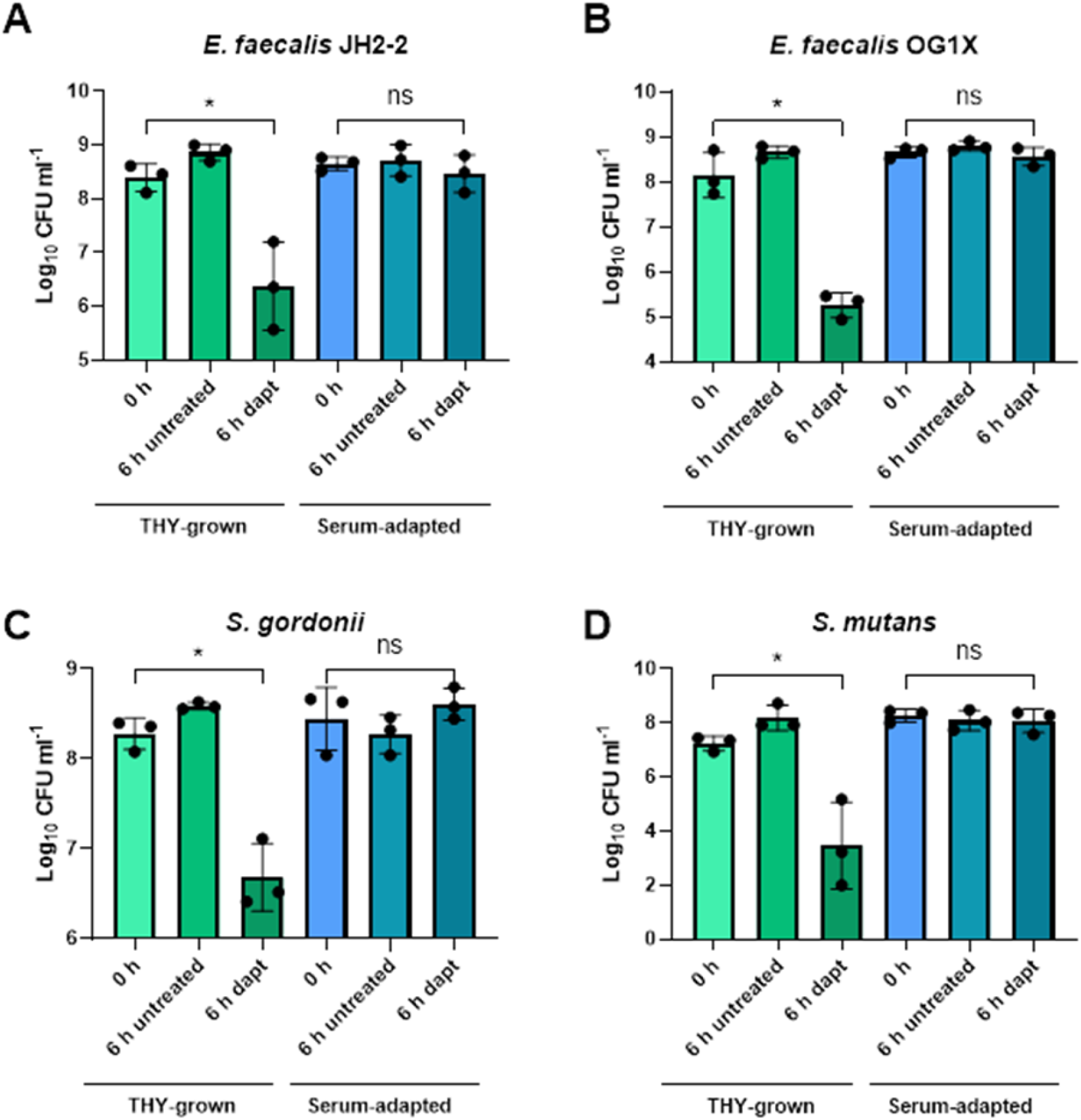
The incubation of *E. faecalis, S. gordonii* and *S. mutans* in human serum induces tolerance to daptomycin. *E. faecalis* JH2-2 **(A)**, *E. faecalis* OG1X **(B)**, *S. gordonii* DL1 **(C)** and *S. mutans* UA159 **(D)** were pre-incubated for 0 (THY) or 16 (Serum) h in serum. Data demonstrate log_10_ CFU ml^-1^ survival after 6 h of exposure to 0 or 80 μg ml^-1^ daptomycin. Graphs represent the geometric mean of three independent experiments and error bars represent the geometric standard deviation of the mean. Data were analysed by one-way ANOVA with Tukey’s multiple comparisons test. *p = < 0.05, ns = not significant.

### The incubation of *E. faecalis, S. gordonii* and *S. mutans* in human serum results in peptidoglycan accumulation

Having established that the incubation of bacteria in serum induced tolerance to daptomycin, the next step was to determine the mechanism responsible. Since our previous work found a role for peptidoglycan accumulation in serum-induced daptomycin tolerance of *S. aureus* [28], we tested whether a similar mechanism was responsible in *E. faecalis* and VGS.

To test this hypothesis, HADA, a blue fluorescent D-alanine analogue that is incorporated into peptidoglycan as it is synthesised, was added during the generation of exponential-phase and serum-adapted bacteria to measure and compare levels of cell wall synthesis [34]. For each strain, bacteria incubated in serum contained significantly more HADA incorporated into the cell wall than exponential-phase cells (Fig. 2A-D), as measured by the fluorescence intensity of individual cells by microscopy. These findings indicated that bacteria accumulated peptidoglycan during incubation in serum, resulting in a thicker cell wall than that seen in broth-grown bacteria.

**Figure 2.**
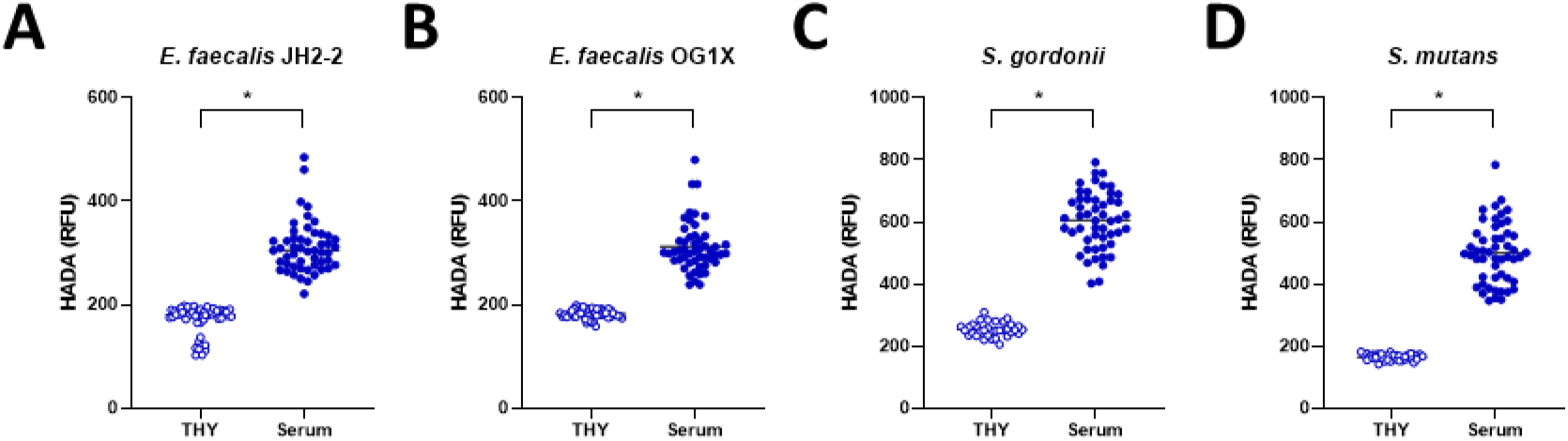
The incubation of *E. faecalis, S. gordonii* and *S. mutans* in human serum triggers accumulation of peptidoglycan. The fluorescence intensity of 50 individual cells of *E. faecalis* JH2-2 **(A)**, *E. faecalis* OG1X **(B)**, *S. gordonii* DL1 **(C)** and *S. mutans* UA159 **(D)** due to incorporation of HADA into the cell wall. Bacteria were either examined in exponential phase (THY) or after incubation in serum (Serum). Graphs show measurements of individual cells with the median indicated and were analysed by paired t-test. *p = <0.05.

### Inhibition of peptidoglycan accumulation partially blocks serum-induced daptomycin tolerance

To test whether serum-induced cell wall accumulation contributed to daptomycin tolerance, peptidoglycan biosynthesis was inhibited during bacterial incubation in serum using the antibiotic fosfomycin, which targets MurA, before subsequent challenge with the lipopeptide antibiotic [38].

Using a range of fosfomycin concentrations, we found that the presence of this antibiotic in serum reduced the development of daptomycin tolerance in all four strains at concentrations that did not directly kill bacteria (Fig. 3A,B,C,D). In the case of JH2-2, bacteria incubated in serum without fosfomycin were highly tolerant of daptomycin, but the presence of the MurA inhibitor at the highest concentrations (80 or 320 µg ml^-1^) reduced bacterial survival ∼90 % after exposure to the lipopeptide antibiotic (Fig. 3A). Similar findings occurred with *E. faecalis* OG1X at the highest fosfomycin concentration, with > 90 % reduction in tolerance compared to bacteria incubated in serum alone (Fig. 3B). Serum-induced daptomycin tolerance of *S. gordonii* was also reduced ∼90 % by the presence of fosfomycin at 80 µg ml^-1^, but this bacterium was killed by the highest concentration of the peptidoglycan synthesis inhibitor, obscuring any impact on daptomycin tolerance (Fig. 3C). Finally, fosfomycin inhibited daptomycin tolerance of *S. mutans* > 90 % at the highest concentration (Fig. 3D). Taken together, these data confirmed our earlier finding that serum induces a high level of daptomycin tolerance and provided evidence that this phenotype is partially due to the accumulation of peptidoglycan.

**Figure 3.**
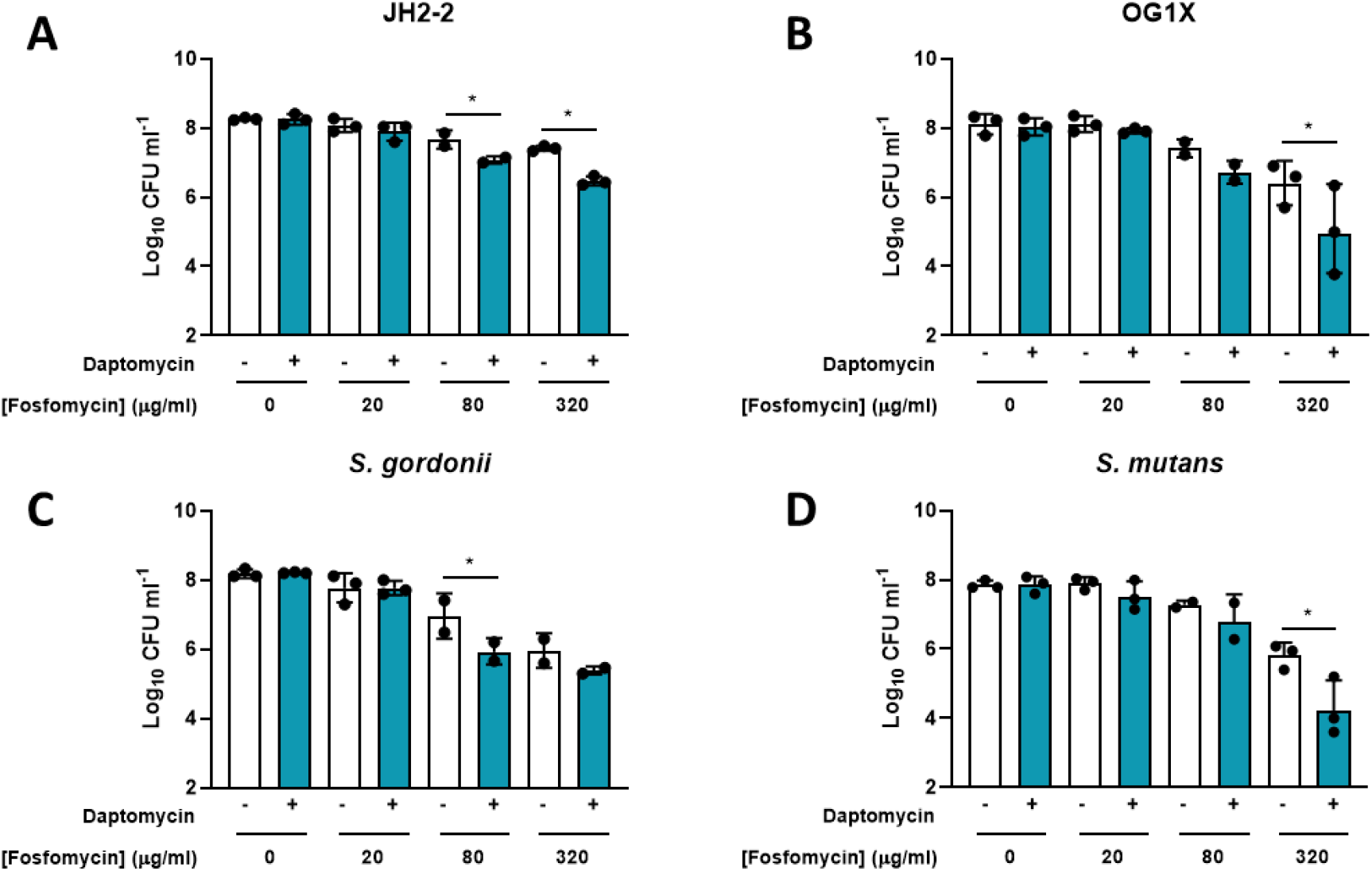
Inhibition of peptidoglycan synthesis in bacterium in serum reduces the development of daptomycin tolerance. *E. faecalis* JH2-2 **(A)**, *E. faecalis* OG1X **(B)**, *S. gordonii* DL1 **(C)** and *S. mutans* UA159 **(D)** were incubated in serum without or with the indicated concentrations of fosfomycin before exposure to daptomycin (80 μg ml^-1^) or not for 6 h and survival measured by determination of CFU counts. Graphs in **A-D** show the geometric mean from three independent experiments and error bars represent the geometric standard deviation of the mean. Data were analysed by one-way ANOVA with Sidak’s multiple comparisons test. *p < 0.05.

### Cardiolipin synthases contribute to serum-induced daptomycin tolerance in *E. faecalis*

In addition to peptidoglycan accumulation, serum-induced daptomycin tolerance in *S. aureus* occurs via an increase in cardiolipin abundance in the membrane. Therefore, we examined serum-induced daptomycin tolerance in *E. faecalis* mutant strains defective for one or both of the cardiolipin synthases (Δ*cls1*, Δ*cls2*) and/or one of the two lysylphosphatidylglycerol synthases present in *E. faecalis* (Δ*mprF2*) [39].

Mutant strains lacking either one of the cardiolipin synthases or the lysylphosphatidylglycerol synthase MprF2 were similarly susceptible to daptomycin as the wild type, with efficient killing of broth-grown bacteria and high-level tolerance of serum-adapted cells (Fig. 4A,B). However, a mutant lacking both cardiolipin synthase genes was significantly less tolerant than wild type of daptomycin after incubation in serum, with ∼4-fold fewer bacteria surviving exposure to the lipopeptide antibiotic (Fig. 4B).

**Figure 4.**
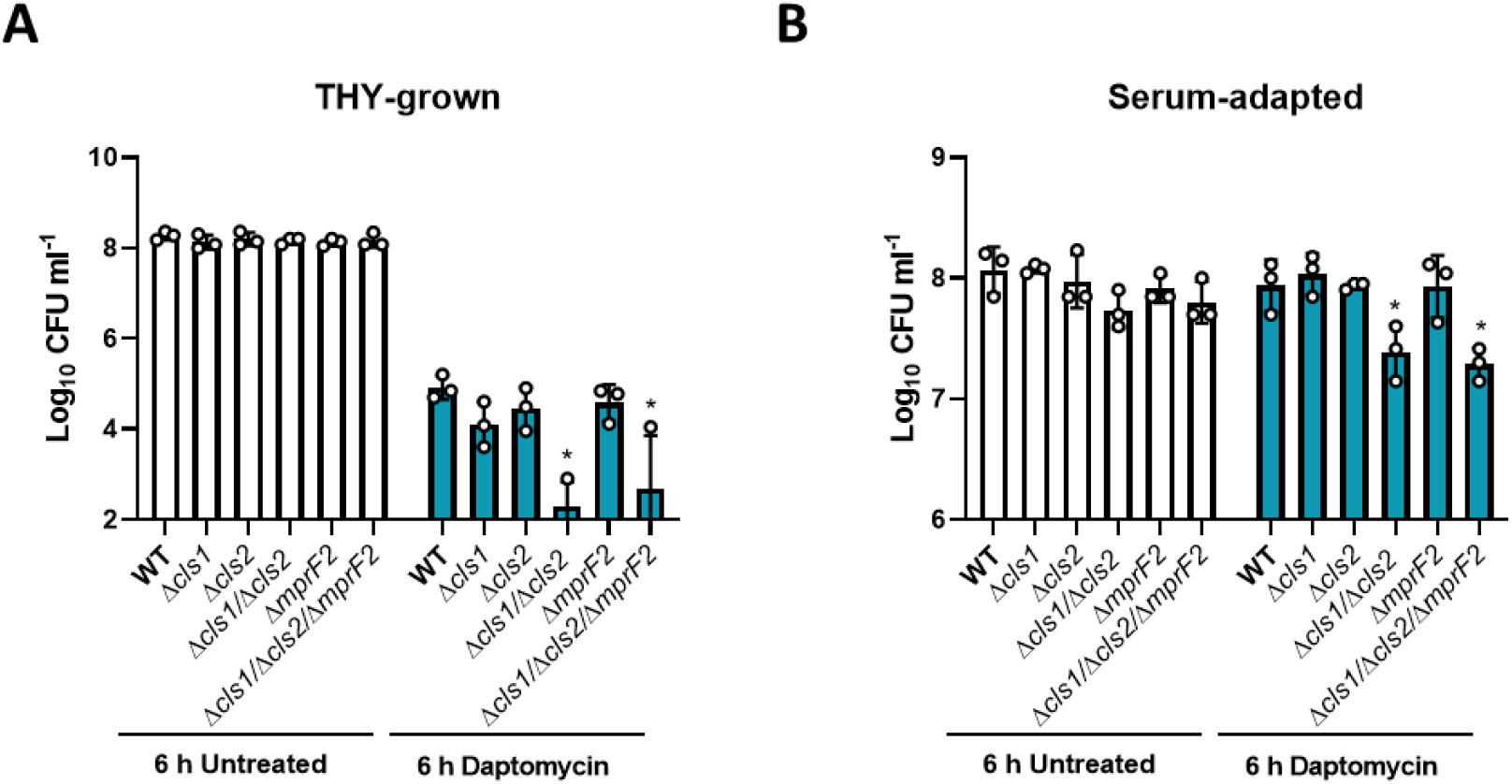
Cardiolipin synthases contribute to serum-induced daptomycin tolerance. THY-grown (**A**) and serum-adapted (**B**) cultures of *E. faecalis* OG1RF WT, Δ*cls1*, Δ*cls2*, Δ*cls1/*Δ*cls2*, Δ*mprF2*, Δ*cls1/*Δ*cls2/*Δ*mprF2* were exposed, or not, to 80 µg ml^-1^ daptomycin for 6 h and survival determined by CFU counts. Data represent the geometric mean ± geometric standard deviation of three independent experiments. Data were analysed by two-way ANOVA with Dunnett’s multiple comparisons test. * p < 0.05, WT vs mutant.

The additional absence of *mprF2* had no effect on survival, suggesting that cardiolipin but not lysylphosphatidylglycerol contributes to serum-induced daptomycin tolerance, as occurs in *S. aureus*. However, the double Δ*cls1/*Δ*cls2* mutant was also more susceptible to killing by daptomycin when in exponential phase and so it is not clear to what extend cardiolipin synthesis contributes to the serum-induced daptomycin tolerance phenotype (Fig. 4A).

### The contribution of cardiolipin synthases to tolerance is independent of peptidoglycan accumulation

Finally, we aimed to determine whether synthesis of peptidoglycan and cardiolipin were sufficient to explain the serum-induced daptomycin tolerance phenotype in *E. faecalis*. To do this, we incubated the wild type and Δ*cls1*/Δ*cls2* double mutant in serum with or without fosfomycin to inhibit peptidoglycan synthesis and then measured bacterial survival after daptomycin exposure.

As expected from previous experiments, fosfomycin treatment reduced serum-induced tolerance in a dose-dependent manner, whilst bacteria defective for both cardiolipin synthases were also less tolerant than the wild type (Fig. 5). However, when the Δ*cls1*/Δ*cls2* was incubated in serum in the presence of fosfomycin, there was a further decrease in tolerance beyond what was seen in the absence of the cell wall synthesis inhibitor (Fig. 5). Therefore, maximal serum-induced daptomycin tolerance requires synthesis of peptidoglycan and cardiolipin.

**Figure 5.**
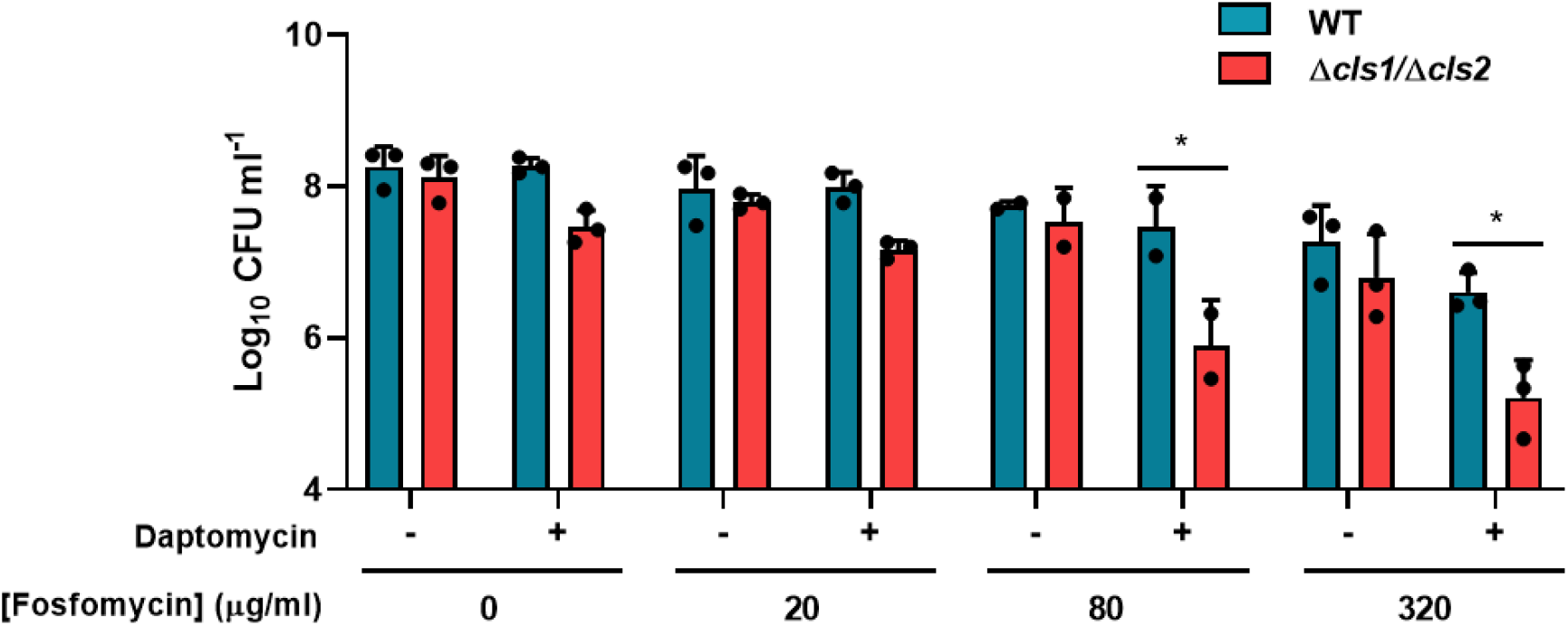
Peptidoglycan and cardiolipin synthesis contribute to serum-induced daptomycin tolerance. *E. faecalis* OG1RF WT and Δ*cls1/*Δ*cls2* were incubated in serum without or with the indicated concentrations of fosfomycin before exposure to daptomycin (80 μg ml^-1^) or not for 6 h and survival measured by determination of CFU counts. Data represent the geometric mean ± geometric standard deviation of three independent experiments. Data were analysed by two-way ANOVA with Sidak’s multiple comparisons test. *p < 0.05.

## Discussion

The key finding from this work is that representative strains of IE pathogens *E. faecalis* and VGS acquire daptomycin tolerance after incubation in human serum, in part because of peptidoglycan accumulation. We also found that cardiolipin synthesis contributes to serum-induced daptomycin tolerance in *E. faecalis*, although this may reflect a general increase in susceptibility to the lipopeptide antibiotic due to the lack of cardiolipin in the membrane.

These findings are consistent with *S. aureus*, which also acquires a very high degree of daptomycin tolerance after incubation in serum due to increased abundance of cardiolipin and peptidoglycan [28]. However, whilst changes to the cell wall and membrane explain serum-induced daptomycin tolerance in *E. faecalis*, an additional mechanism has been reported [40,41,42]. Incubation of *E. faecalis* in serum leads to changes in membrane fluidity that confer reduced susceptibility to killing by the lipopeptide antibiotic, independently of induction of a membrane stress response [40,41,42].

The finding that a thickened cell wall reduces daptomycin susceptibility is similar to observations made with vancomycin-intermediate *S. aureus* (VISA) strains, which are also less susceptible to the lipopeptide antibiotic [43].

The partial inhibition of tolerance induction by fosfomycin indicate that these two antibiotics could make an effective combination therapy for treating enterococcal and streptococcal IE. Synergy between daptomycin and fosfomycin against *S. aureus* has been previously investigated *in vitro* and in *in vivo* animal models [44,45]. For example, García-de-la-Mària *et al*. demonstrated that, in comparison to daptomycin monotherapy, combination therapy with fosfomycin significantly increased the proportion of sterile vegetations (72% versus 100%) and decreased the density of bacteria in the remaining vegetations in an experimental rabbit model of methicillin-resistant *S. aureus* (MRSA) IE [45]. There is also evidence that daptomycin-fosfomycin is safe and effective in the treatment of staphylococcal IE in humans, although this report only examined three cases [46]. Furthermore, treatment of patients with MRSA bacteraemia with daptomycin-fosfomycin combination therapy was associated with lower rates of treatment failure than daptomycin alone [46]. However, combination therapy did not improve patient survival and was associated with nephrotoxicity [46]. Therefore, additional work is needed to understand how best to use fosfomycin with daptomycin to maximise patient benefit and there is also a paucity of evidence for the use of fosfomycin in combination with daptomycin to treat IE caused by enterococci or streptococci.

In summary, this study has shown that bacterial incubation in serum results in cell wall thickening, which confers tolerance to daptomycin, presumably because the lipopeptide antibiotic is too large to traverse a thickened cell wall. However, cell wall thickening can be blocked using fosfomycin, offering the possibility of new treatment approaches with enhanced efficacy.

## Acknowledgements

Angela Nobbs (University of Bristol), Stéphane Mesnage (University of Sheffield) and Elizabeth Fozo (The University of Tennessee Knoxville) are gratefully acknowledged for providing strains.

## Author contributions

EVKL and AME conceived the project. ARHT & EVKL conducted experiments and analysed data. EVKL and AME directed the project. All authors contributed to writing the manuscript.

